# A UL26-PIAS1 complex antagonizes anti-viral gene expression during Human Cytomegalovirus infection

**DOI:** 10.1101/2024.02.19.580933

**Authors:** Jessica Ciesla, Kai-Lieh Huang, Eric J. Wagner, Joshua Munger

**Affiliations:** Department of Biochemistry and Biophysics, University of Rochester School of Medicine and Dentistry, Rochester, New York, USA

**Author notes:** Corresponding addressed to Joshua Munger.

## Abstract

Viral disruption of innate immune signaling is a critical determinant of productive infection. The Human Cytomegalovirus (HCMV) UL26 protein prevents anti-viral gene expression during infection, yet the mechanisms involved are unclear. We used TurboID-driven proximity proteomics to identify putative UL26 interacting proteins during infection to address this issue. We find that UL26 forms a complex with several immuno-regulatory proteins, including several STAT family members and various PIAS proteins, a family of E3 SUMO ligases. Our results indicate that UL26 prevents STAT phosphorylation during infection and antagonizes transcriptional activation induced by either interferon α (IFNA) or tumor necrosis factor α (TNFα). Additionally, we find that the inactivation of PIAS1 sensitizes cells to inflammatory stimulation, resulting in an anti-viral transcriptional environment that mirrors ΔUL26 infection. Further, PIAS1 is important for HCMV cell-to-cell spread, which depends on the presence of UL26, suggesting that the UL26-PIAS1 interaction is vital for modulating intrinsic anti-viral defense.

## Introduction

Human Cytomegalovirus (HCMV) is a widespread pathogen belonging to the *Herpesviridae* family that causes severe morbidity during congenital infection and in immunosuppressed individuals. In the United States, 1 in 200 children are born with HCMV, and of this population, approximately 10% are born with symptoms including blindness, deafness, microcephaly, jaundice, seizures, and, in the most severe cases, death (1). Other reports suggest up to 20-25% of HCMV-positive newborns are symptomatic at birth or will develop long-term medical complications (2). Currently, there are no FDA-approved vaccines for HCMV, and anti-viral therapeutic options exhibit toxicity, low bioavailability, and the emergence of drug-resistant strains in immunosuppressed populations (3, 4, 5). Understanding the determinants of successful HCMV infection could provide novel points of therapeutic intervention to lessen the disease burden of HCMV.

The host’s innate immune response is critical for limiting viral infection. Activation of innate immune pathways, including nuclear factor kappa-light-chain-enhancer of activated B cells (NFκB) (6, 7, 8) and Janus kinase/signal transducer and activator of transcription (JAK/STAT) (9, 10) promote anti-viral gene expression to restrict viral infection. Viruses have evolved mechanisms to evade detection or block innate immune activation to replicate efficiently. With respect to HCMV, the virion tegument contains a matrix of proteins between the nucleocapsid and envelope that are released into the cytoplasm immediately upon infection. Many tegument proteins antagonize various cellular anti-viral activities. The most abundant tegument protein, pp65, blocks IRF3 activation to restrict type I interferon signaling (11). The pp71 protein binds the innate immune effector Daxx to prevent Daxx-mediated silencing of the HCMV major Immediate Early Promoter (MIEP) (12). The UL23 protein restricts Type II interferon gene expression during infection by interacting with the human n-Myc interactor (13). The UL26 tegument protein, which is important for high titer replication (14, 15), restricts the expression of cytokine signaling genes during infection (16) and antagonizes TNFα-induced NFκB activation (17). However, the mechanisms by which UL26 modulates host innate immune signaling and supports high titer viral replication are unclear.

Here, we explored the UL26 interactome during early-stage infection using proximity-based proteomics (18). To identify UL26-associated proteins that likely contribute to high titer viral replication, we compared the interactome of wildtype UL26 with that of a UL26 C-terminal deletion mutant (UL26ΔC) that phenocopies UL26-deletion viral growth kinetics (19). Our data suggest that the full-length UL26 protein, but not UL26ΔC, interacts with PIAS1 (protein inhibitor of activated STAT 1), a key modulator of STAT and NFkB transcription factors (20). PIAS proteins negatively regulate anti-viral gene expression by binding DNA at specific promoter regions and antagonizing transcription factor binding (21, 22). PIAS proteins also act as E3-SUMO ligases and conjugate a small ubiquitin-modifier (SUMO) polypeptide to modulate protein activity, including inhibition of transcription factor activity (23). Our results indicate that cells lacking PIAS1 are less permissive to HCMV spread and have increased expression of anti-viral genes, similar to what is observed during ΔUL26 infection.

## Results

### The proximal UL26 interactome contains several host-defense regulatory proteins

To investigate the mechanisms through which UL26 contributes to HCMV replication, we sought to identify proteins that interact with UL26 during infection. Towards this end, we fused UL26 to the TurboID biotin ligase, which conjugates biotin to proximal proteins with faster kinetics relative to previous biotin ligase domains (18). In addition to TurboID tagging WT UL26, we also tagged a UL26 variant in which the C-terminal 38 amino acids have been deleted (UL26ΔC). Recombinant HCMV strains that express this UL26ΔC variant behave similarly to ΔUL26 strains with respect to their growth kinetics (19). Given that UL26’s C-terminus is critical for its function (19), we reasoned that the proteins interacting with WT UL26, but not the UL26ΔC mutant, are more likely to be important for UL26’s contributions to HCMV infection. We created recombinant HCMV strains containing TurboID-tagged versions of WT UL26 (Tur-N-UL26wt) or the UL26ΔC mutant (Tur-N-UL26ΔC) in the original UL26 genomic locus (Fig. 1a). Both TurboID-fused proteins were stably expressed, accumulated normally during infection (Fig. 1b), and localized to the nucleus as expected (Fig. 1c). Further, TurboID tagging of WT UL26 did not impact HCMV viral titers, suggesting that UL26 was still functional upon tagging (Fig. 1d). Similarly, TurboID tagging of UL26ΔC grew to a similar titer as the ΔUL26 virus (Fig. 1d).

**Figure 1.**
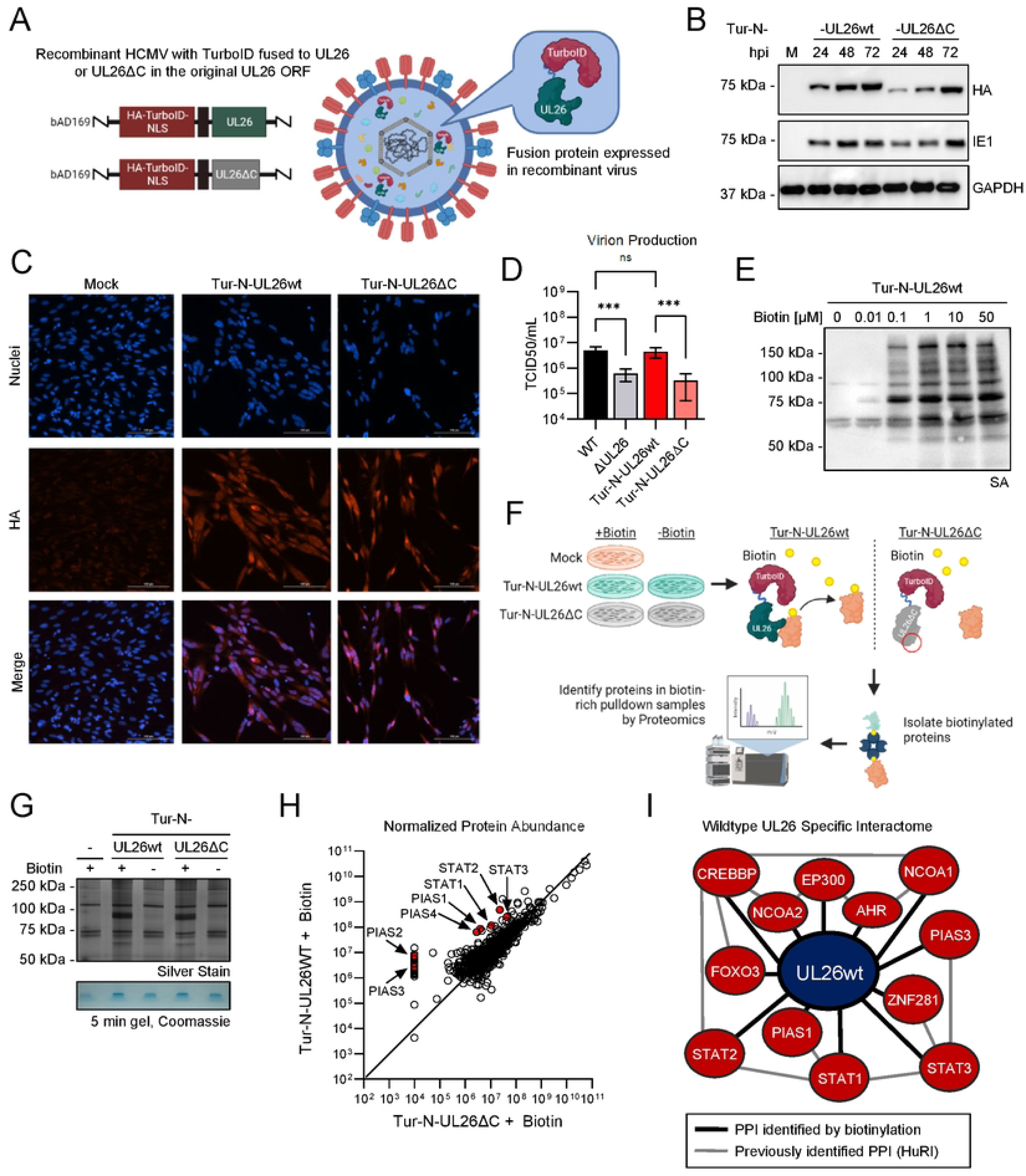
The proximal UL26 interactome contains several host-defense regulatory proteins. **(A)** Recombinant HCMV, strain AD169. containing TurbolD fused to the N-terminus of wildtype UL26 (Tur-N-UL26wt) or UL26ΔC (Tur-N-UL26ΔC) in the UL26 ORF. N-terminus of TurbolD fused to a 3x HA tag. and the C-terminus is fused to a 3x SV40 nuclear localization signal (NLS or N). (B) Western blot showing the expression of TurbolD fused to UL26wt or UL26ΔC in MRC5s infected with Tur-N-UL26wt or Tur-N-UL26ΔC (MOI = 3) at the indicated hour post infection (hpi) compared to mock (M). (C) Immunofluorescence images of MRC5s infected with Tur-N-UL26wt or Tur-N-UL26ΔC (MOI = 3) at 48 hpi. Tur-N-UL26wt and Tur-N-UL26ΔC protein detected by anti-HA: nuclei stained with Hoechst. (D) Infectious virions produced by MRC5s infected with wild-type HCMV (WT). UL26-deletion (ΔUL26). Tur-N-UL26wt or Tur-N-UL26ΔC at 120 hpi. determined by TCID50 analysis. Virion production is represented as the mean of 6 biological replicates +/- SEM. FDR-adjusted p-values determined using 2-way ANOVA followed by two-stage step-up method of Benjamin), Krieger and Yekutieli n.s = not significant. . *p<0.033. **p<0.002, ***p<0.001. (E) Western blotting showing the accumulation of biotinylated protein in MRC5s infected with Tur-N-UL26wt (MOI = 3). Infected cells were incubated with the indicated concentration of biotin at 23 hpi, and protein was harvested at 24 hpi. (F) Overview of TurbolD-mediated proximity labeling during infection with Mock. Tur-N-UL26wt. or Tur-N-UL26ΔC (MOI = 3). Total protein harvested 24 hpi, biotinylated proteins isolated by affinity purification using streptavidin-bound magnetic beads then identified by LC-MS/MS. (G) Silver stain showing total biotin-enriched protein in affinity pulldown samples (top). Remaining affinity pulldown sample concentrated on a 5 minute gel and stained with Coomassie for proteomic analysis (bottom). (H) Protein abundance data from LC-MS/MS of Tur-N-UL26wt and Tur-N-UL26ΔC affinity pulldown samples median-normalized and represented as a scatter plot. STAT and PIAS proteins highlighted in red. (I) Interactome of 68 proteins enriched in Tur-N-UL26wt+Biotin relative to Tur-N-UL26ΔC+Biotin (Fold Change > 5) identified using the Human Reference Protein Interactome Mapping Project (HuRI) database. Black lines indicate wildtype UL26-specific interactions detected in this proximity proteomics experiment, grey lines indicate previously identified protein-protein interactions from HuRI.

We next assessed the biotinylation activity of our TurboID-tagged recombinant viruses as well as the concentration of biotin that would yield robust proximal biotinylation over an hour time frame. Our results indicate pulsing with 1 μM biotin resulted in maximal biotinylation as assessed by streptavidin HRP signal (Fig. 1e). Moving forward to the proteomic analysis, fibroblasts were mock infected or infected with Tur-N-UL26wt, or Tur-N-UL26ΔC (MOI = 3) for 23 hours, treated with 1 μM biotin or vehicle for 1 hour, and purified biotinylated proteins via streptavidin affinity (Fig. 1f). The resulting proteins were analyzed by silver stain (Fig. 1g). As expected, streptavidin affinity purified more proteins in the infected samples treated with biotin relative to either the non-biotin treated infected cells or the mock-infected biotin treated cells (Fig. 1g, top). Purified samples were then concentrated by 5-minute SDS-PAGE, stained for total protein (Fig. 1g, bottom), and analyzed by LC-MS/MS to identify the biotinylated proteins.

In total, LC-MS/MS identified 601 Human and HCMV proteins in the streptavidin affinity purified samples (Table S1). Among those 601 identified proteins, several were preferentially biotinylated by the full-length UL26-TurboID versus UL26ΔC-TurboID (Fig. 1h). Specifically, sixty-eight proteins were ≥5-fold enriched with respect to interacting with WT UL26 relative to the UL26ΔC mutant, suggesting the UL26 C-terminus is necessary for their interaction (Table S2). The UL31 protein was the only viral protein among these 68 proteins and was ∼8-fold enriched in the wildtype UL26 sample relative to the UL26ΔC sample (Table S3). Twenty additional viral proteins were found to interact with both WT UL26 and the UL26ΔC mutant (Table S3). Of the 67 human proteins that interacted preferentially with the full-length UL26, many are involved in innate immune signaling, including several STAT and PIAS family members (Fig. 1h, & Table S2). Further, several of these proteins are thought to be present in common complexes, as predicted by the Human Reference Interactome (HuRI) (24) (Fig. 1i). Collectively, these results suggest that UL26 associates with a complex involved in the modulation of host responses to infection, consistent with activities previously linked to UL26 (17, 25).

### The UL26 protein blocks STAT activation and cytokine-mediated transcriptional activation

While UL26 has been found to attenuate NFkB signaling (17), the associated proteins identified above suggest it may also modulate STAT signaling. To explore this possibility, we analyzed STAT phosphorylation in WT HCMV or UL26ΔC infected cells. Phospho-STAT3 levels were significantly increased in UL26ΔC-infected cells relative to mock or WT HCMV infection (Fig. 2a). Similarly, phospho-STAT1 and phospho-STAT2 levels were increased in UL26ΔC-infected cells relative to mock or WT infection (Fig. 2b-c). These results suggest that the UL26 C-terminus is necessary to restrict STAT phosphorylation during HCMV infection.

**Figure 2.**
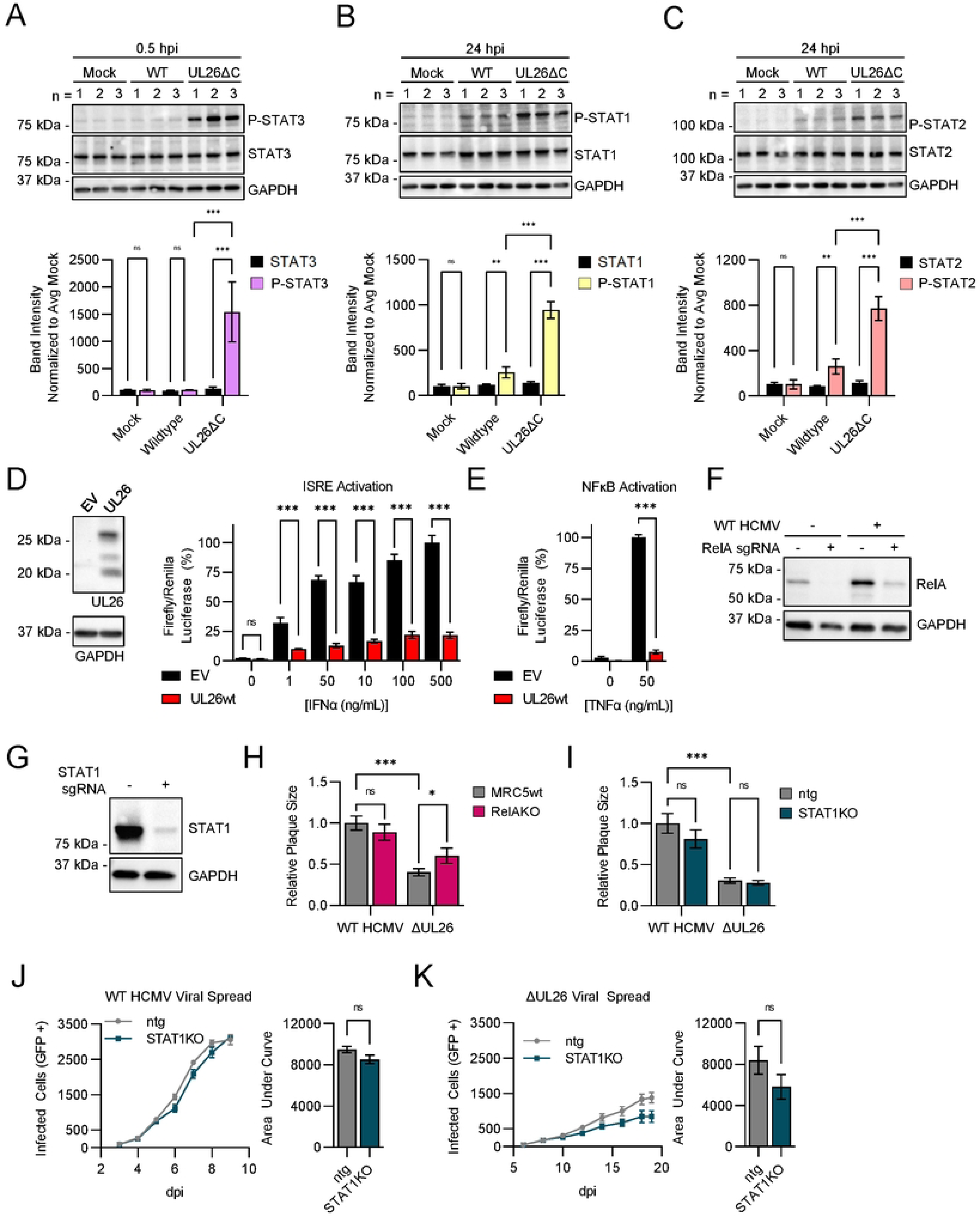
The UL26 protein blocks TNFα and IFNα signaling. (A-C) MRC5 cells infected with mock, wildtype HCMV, or UL26ΔC mutant virus (MOI = 3). Protein harvested from three independent biological replicates at (A) 0.5 or (B and C) 24 hpi and processed by western blot as indicated (top). For each replicate, band intensity for the indicated protein was quantified relative to GAPDH and normalized to the average mock-infected band intensity (100). Data is represented as the mean value of three biological replicates+/- SD (bottom). FDR-adjusted p-values determined using 2-way ANOVA followed by two-stage step-up method of Benjamini. Krieger and Yekutieli n.s = not significant. *p<0.033, **p<0.002, ***p<0.001. (D and E) Dual-Glo luciferase assay in HEK293T cells containing a renilla luciferase plasmid and Interferon Stimulated Response Element (ISRE) firefly luciferase reporter plasmid (D). or NFkB firefly luciferase reporter plasmid (E). UL26 is expressed in HEK293T cells transfected with a UL26 expression plasmid for 48 hr (D, left). HEK293T cells containing Dual-Glo luciferase reporter assay system were transfected with empty vector (EV) or a UL26 expression plasmid for 48 hr, then stimulated with the indicated concentration of IFNα (D) or TNFα (E) for 24 hr. Firefly andrenilla luciferin were quantified using a DualGlo luciferase kit. For each sample, firefly luciferin was normalized to renilla luciferin and represented as the mean of 4 biological replicates +/- SE. FDR-adjusted p-values determined using 2-way ANOVA followed by two-stage step-up method of Benjamini. Krieger and Yekutieli n.s = not significant, *p<0.033, **p<0.002, ***p<0.001. (F) Immunoblot of protein harvested from MRC5s transfected with CRISPR-Cas9 complexed to RelA-targeted sgRNA infected with mock or wildtype HCMV. (G) Immunoblot of protein harvested from confluent MRC5 cells transfected with CRISPR-Cas9 complexed to STAT1-targeted sgRNA or a non-targeting guide (ntg). (H and I) Plaque size of wildtype HCMV or ΔUL26 in RelAKO cellscompared to parental MRC5s (H) or STAT1KO cells compared to ntg cells (I). The area of 20 representative plaques were quantified for each condition and normalized to the mean plaque size of wildtype HCMV in parental (H) or ntg (I). Mean plaque size for each condition is represented as the mean +/- SE. FDR-adjusted p-values determined using 2-way ANOVA followed by two-stage step-up method of Benjamini, Krieger and Yekutieli n.s = not significant, *p<0.033, **p<0.002, ****p<0.001. (J and K) Viral spread assay of GFP-expressing wildtype HCMV (J, MOI = 0.01) or GFP-expressing ΔUL26 virus (K, MOI = 0.01) in STAT1KO (teal) cells compared to ntg (grey). Number of infected cells (GFP+) quantified for 6 biological replicates at the indicated day post infection (dpi) and represented as the mean +/- SE in an XY-plot (left) for each day. Total area under curve (AUC) calculated for viral spread in each cell line and plot as a bargraph +/- SE (right); t-values determined by student’s unpaired t-test, ns = not significant, *t<0.033, **t<0.002, ***t<0.001.

We previously have found that UL26 is sufficient to antagonize TNFα-mediated NFκB activation (17) and is necessary to prevent the induction of interferon-stimulated gene (ISG) expression during infection (16). Given our findings that STAT phosphorylation is induced in UL26ΔC infection, we assessed whether UL26 is sufficient to inhibit activation of the Interferon Stimulated Response Element (ISRE), a promoter for ISGs induced by activated STAT transcription factor complexes (26). HEK293T cells were transfected with an ISRE firefly luciferase reporter plasmid and control vector (EV) or UL26-expression vector (Fig. 2d, left). Treatment with increasing amounts of IFNα induced ISRE-dependent luciferase activity, which was inhibited by UL26 expression (Fig. 2d, right), suggesting that UL26 is sufficient to antagonize IFNα-induced ISRE activation. We also analyzed the impact of UL26 expression on TNFα- induced NFkB activation and found that similar to previous results (17) UL26 expression was sufficient to inhibit TNFα-induced activation of NFkB-dependent luciferase activity (Fig. 2e). Together, these data indicate that UL26 is sufficient to antagonize IFNα and TNFα-induced transcriptional activation.

Given that ΔUL26 infection fails to block the expression of cytokine signaling genes (16), we assessed whether the inactivation of STAT1 or RELA, endpoint transcription factors associated with activated STAT and NFkB signaling, respectively, would rescue ΔUL26 replication defects. RelA and STAT1 knockout (KO) MRC5 cell lines were generated through CRISPR-CAS9 gene targeting, resulting in a significant knockdown in expression in polyclonal cell populations relative to non-targeting guide controls (ntg) (Fig. 2f & g). RELA targeting increased the plaque size associated with ΔUL26 infection, although these plaques were still substantially smaller than WT (Fig. 2h). In contrast, the absence of STAT1 did not impact ΔUL26 plaque size (Fig. 2i). Further, the viral spread of WT HCMV or ΔUL26 was not impacted by the lack of STAT1 (Fig. 2j & k). These results suggest that inactivation of either RELA or STAT1 alone is not sufficient to rescue the viral spread defects associated with ΔUL26 infection.

### PIAS protein abundance and SUMOylation are enhanced during HCMV infection

To gain further clues about the possible activities of the UL26-proximal complex identified above, we performed an ontology analysis of the 67 Human proteins enriched in the WT UL26 versus UL26ΔC samples (Table S4) (24). Our analysis revealed that these proteins were preferentially associated with various aspects of protein SUMOylation (Fig. 3a). SUMO proteins are small (∼10 kDa) protein modifiers that are covalently attached to target proteins by E3-SUMO ligases, including Protein Inhibitor of Activated STAT (PIAS) proteins, in a process known as SUMOylation. SUMOylation is a reversible post-translational modification that controls diverse molecular functions, including chromatin structure, DNA repair, transcription, cell cycle progression, and the innate immune response (27). SUMOylation supports the activation of intrinsic immunity, and SUMOylation of some viral proteins restricts viral replication, yet many viruses have evolved mechanisms to modulate SUMOylation and counter this anti-viral activity (28, 29, 30). For HCMV, SUMO over-expression has been found to enhance HCMV replication (31), suggesting that it might benefit infection. Consistent with these findings, we observe protein SUMOylation is enhanced during WT HCMV infection (Fig. 3b). WT HCMV infection also substantially increased the accumulation of PIAS-type E3-SUMO ligases, including PIAS1, PIAS3, and PIAS4, as well as PIAS2, albeit to a much lesser extent (Fig. 3c). Similar results were observed during UL26ΔC infection, SUMO1 and PIAS1 levels were increased (Fig. 3d). To determine if SUMOylation is broadly important for HCMV replication, we treated cells with the SUMOylation inhibitor Subasumstat (32). Subasumstat treatment resulted in a dose-dependent decrease in HCMV-induced SUMOylation and also inhibited viral spread (Fig. 3e). Further, Sabasumstat treatment attenuated the accumulation of infectious virions by ∼100-fold (Fig. 3g), indicating that SUMOylation is important for HCMV infection.

**Figure 3.**
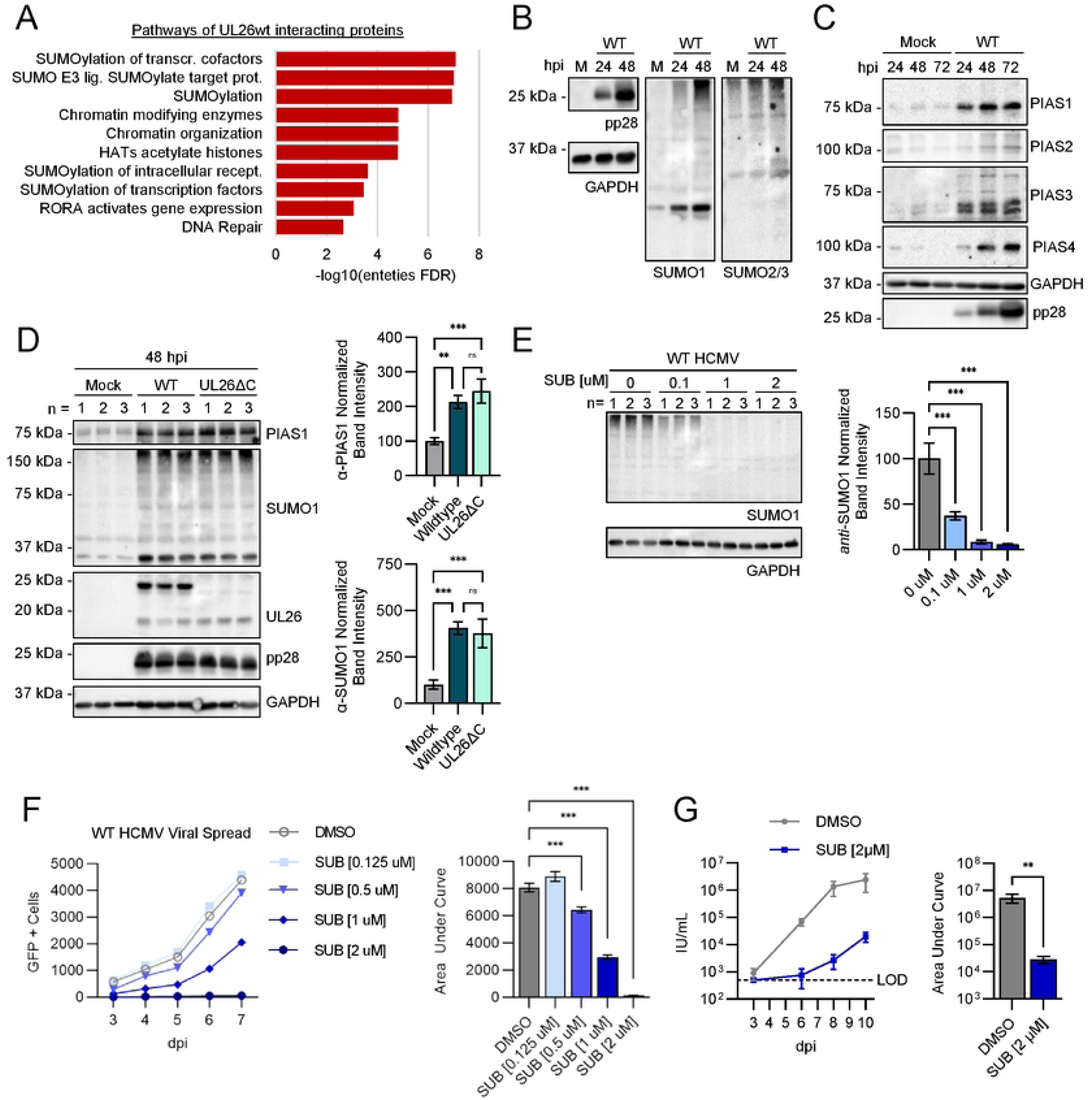
HCMV induces the expression of PIAS family E3 SUMO ligases and increases protein SUMOylation, which is important for HCMV spread. (A) Reactome analysis of 68 proteins enriched in Tur-N-UL26wt+Biotin relative to Tur-N-UL26ΔC+Biotin (Fold Change > 5) Reactome pathways with the top 10 entities FDR represented as -Iog10(entities FDR). (B and C) MRC5 cells infected with mock or wildtype HCMV (WT. MOI = 3). At the indicated hour post infection (hpi), protein harvested for western blot analysis. (D) MRC5 cells infected with mock, WT, or UL26ΔC mutant virus (MOI = 3). Protein harvested at 48 hpi and processed by immunoblot as indicated. Three independent replicates were carried out for each sample group (n = 1-3). For each replicate, band intensity for PIAS1 or SUMO1 was quantified relative to the GAPDH loading control and normalized to the average mock-infected band intensity (100). The mean value of three biological replicates +/- SD were plotted as bar graphs for PIAS1 and SUMO1 (right). FDR-adjusted p-values determined using 2-way ANOVA followed by two-stage step-up method of Benjamini, Krieger and Yekutieli n.s = not significant, *p<0.033, **p<0 002, ***p<0.001. (E) MRC5 cells infected with WT (MOI = 3) and treated with the indicated concentration of Subasumstat (SUB). At 48 hpi, protein harvested and analyzed by western blot (left). Three independent replicates were carried out for each sample group (n = 1-3). For each replicate, band intensity for SUMO1 was quantified relative to the GAPDH loading control and normalized to the average mock-infected band intensity (100). The mean value of three biological replicates +/- SD were plotted as bar graphs (right). FDR-adjusted p-values determined using 2-way ANOVA followed by two-stage step-up method of Benjamini, Krieger and Yekutieli n.s = not significant, *p<0.033, **p<0.002, ***p<0.001. (F) Viral spread assay in MRC5 cells treated with the indicated concentration of Subasumstat (SUB) during infection. Cells infected with GFP-expressing wildtype HCMV (MOI = 0.1). Number of infected cells (GFP + Cells) quantified for 6 biological replicates at the indicated day post infection (dpi) and represented as the mean +/- SE in an XY-plot for each day (left). Total area under curve (AUC) calculated for viral spread in each cell line and plot as a bar graph +/- SE (right); FDR-adjusted p-values determined using 2-way ANOVA followed by two-stage step-up method of Benjamini, Krieger and Yekutieli n.s = not significant, *p<0.033, **p<0.002, ***p<0.001. (G) Viral growth assay of MRC5 cells treated with Subasumstat (2 µM) or DMSO during infection with HCMV (MOI = 0.01). Medium was harvested from cells infected at the indicated dpi. Infectious units (IU) per mL was quantified for each sample and plotted as the mean +/- SEM (left) of 12 biological replicates. Total area under the curve for each treatment was calculated and plotted as a bar graph +/+- SEM (right), t-values determined by student’s unpaired t-test, ns = not significant, *t<0.033, *t<0.002, —t<0.001.

### PIAS1 inactivation potentiates anti-viral gene expression

Our proteomics data indicate that WT UL26 preferentially biotinylated several PIAS family members relative to the UL26ΔC mutant. While each of the different PIAS family members could play important roles during HCMV infection, we focused on PIAS1 as it accumulates to abundant levels at early times post-infection (Fig. 3c). Further, similar to UL26, PIAS1 has been found to modulate multiple innate immune signaling pathways, including SUMOylating STAT1’s C-terminal region, attenuating STAT1’s DNA-binding activity, and inhibiting inflammatory gene expression (33). Additionally, PIAS1 SUMOylates NFkB signaling components, e.g., inhibiting anti-viral gene expression through SUMOylation of IKKα and binding RELA (34, 35). To examine PIAS1’s anti-viral activity, we targeted it via delivery of CRISPR-ribonucleoprotein (RNP) complexes, which substantially reduced PIAS1 protein accumulation in polyclonal cell populations (Fig. 4a). Subsequent RNA-seq analysis of control versus these PIAS1-targeted cells revealed 302 genes significantly more abundant in cells lacking PIAS1 (PIAS1KO) relative to the non-targeting guide (ntg) control cell line (Table S5). Ontology analysis of these 302 genes revealed that PIAS1 inactivation induced genes involved in cholesterol metabolism (Fig. 4b, Table S5), including 3-hydroxy-3-methylglutaryl-CoA reductase (HMGCR), lanosterol synthase (LSS), and mevalonate diphosphate decarboxylase (MVD) (Fig. 4c, Table S5). These findings are consistent with previous reports indicating that PIAS1 suppresses the expression of cholesterol-associated enzymes (36, 37). In addition to cholesterol metabolism, PIAS1 inactivation increased the expression of genes associated with IL12-induced JAK-STAT signaling (Fig. 4b, Table S5).

**Figure 4.**
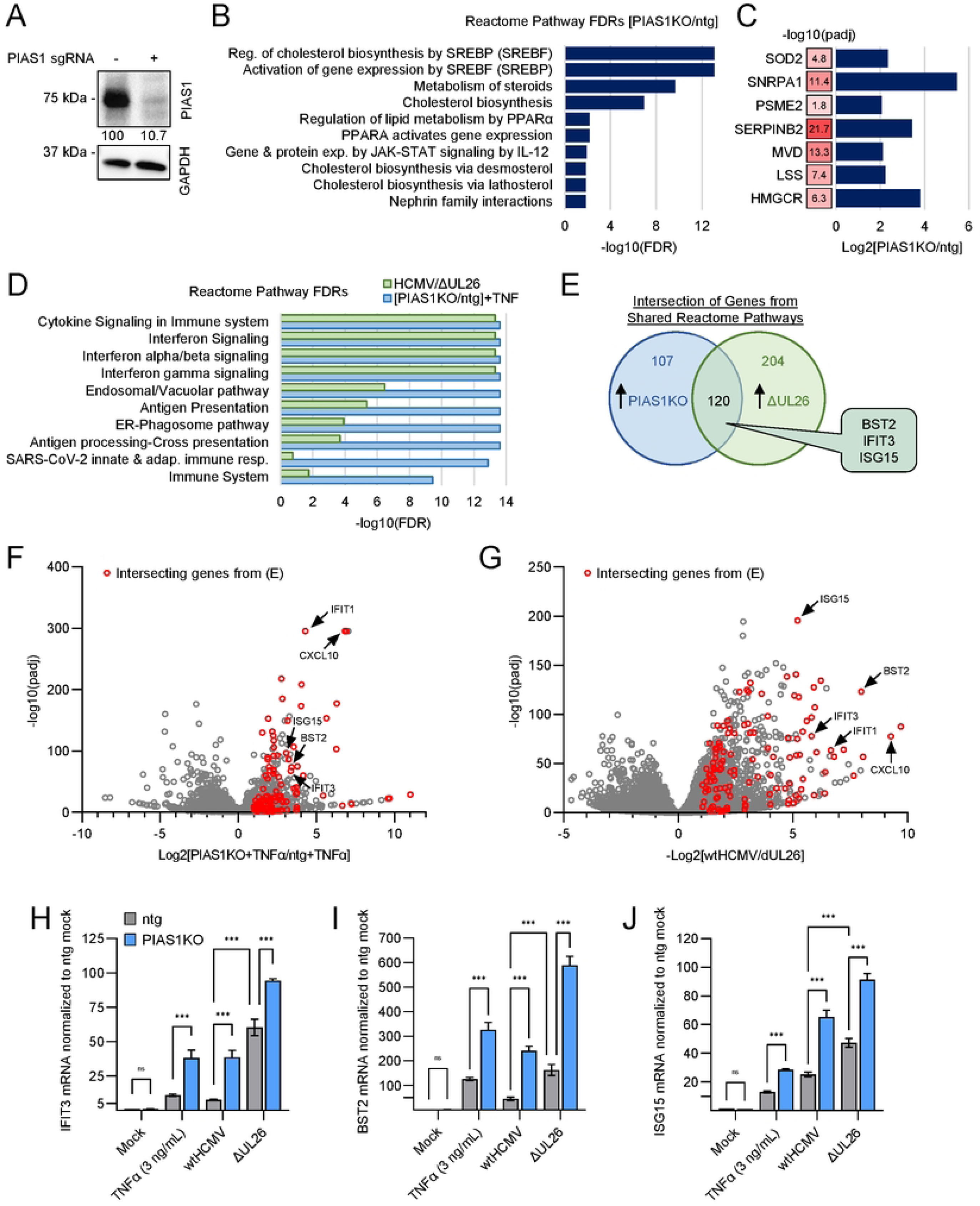
PIAS1 inactivation sensitizes cells to inflammatory stimulation resulting in increased anti-viral gene expression that phenocopies ΔUL26 infection. (A) Western blot of protein harvested from confluent MRC5 cells transfected with CRISPR-cas9 RNP complexed to sgRNAs towards PIAS1 or a non­targeting guide (ntg) RNA Band intensity of PIAS1 normalized to GAPDH for each sample, ntg PIAS1 band intensity standardized to 100. (B - F) Ntg and PIAS1 knockout (KO) cells treated with vehicle or TNFα (3 ng/mL) for 24 hr. RNA harvested and analyzed by RNA-seq. (B) Reactome analysis of genes enriched in vehicle-treated PIAS1KO cells relative to ntg (Log2[PIAS1KO/ntg]> 2). Reactome pathways with the top 10 entities FDR represented as -Iog10(entities FDR). (C) Log2 fold change enrichment in PIAS1KO cells relative to ntg and padj values for individual genes significantly enriched in vehicle-treated PIAS1KO cells relative to ntg. (D, blue) Reactome analysis of genes enriched in TNFα-treated PIAS1KO cells relative to ntg. Reactome pathways with the top 10 entities FDR represented as -Iog10(entities FDR). (D and E, green; F) MRC5 cells infected with WT or ΔUL26 HCMV at MOI = 3 for 24 hr. RNA harvested and analyzed by RNA-seq. (D, green) Reactome pathways of genes enriched in cells infected with ΔUL26 relative to WT that overlap with PIAS1KO RNA seq experiment. Entities FDR is represented as -Iog10(entities FDR). (E) Venn diagram of entities from reactome analysis. Genes enriched in TNFα-treated PIAS1KO cells relative to ntg cells (blue, 227 genes) and ΔUL26 relative to WT (green, 324 genes). (F and G) Volcano plot of (F) RNA-seq data from TNFα-treated PIAS1KO and ntg cells (G) RNA-seq data from cells infected with WT or ΔUL26. 120 intersecting genes shown in E are represented in red. (H - J) RTqPCR quantification of IFIT3 (E) BST2 (F) and ISG15 (G) in ntg and PIAS1KO cells treat with mock, wtHCMV (MOI = 3), ΔUL26 (MOI = 3) or TNFα (3 ng/mL) for 24 hr. Data represented as the mean of 3 biological replicated +/- SD; FDR-adjusted p-values determined using 2-way ANOVA followed by two-stage step-up method of Benjamini, Krieger and Yekutieli n.s = not significant, *p<0.05, **p<0.01, ***p<0.001.

Given that PIAS1 modulates NFkB signaling (34, 35), we tested whether inactivation of PIAS1 impacted the response to TNFα treatment, an activator of NFkB signaling. TNFα-treated PIAS1KO cells exhibited substantially larger increases in genes involved in cytokine signaling and the innate immune response relative to TNFα-treated control cells (Fig. 4d, Table S6 & S7). Notably, similar pathways are induced when comparing ΔUL26 infection relative to WT HCMV infection (Fig. 4d, Table S8, (25)). Of the genes within these intersecting pathways, 120 were significantly induced by both ΔUL26 infection and TNFα-treated PIAS1 KO cells relative to their respective control conditions (Fig. 4e & Table S9). Many of these 120 overlapping genes are interferon-stimulated genes (ISGs) including IFIT1-3, ISG15, CXCL10, and BST2. These genes were among the most strongly induced in both conditions (Fig. 4f & g, red). Collectively, these results suggest that the transcriptional environment induced by ΔUL26 infection mirrors that of TNFα-treated PIAS1KO cells.

Given that PIAS1KO cells were sensitized to TNFα treatment, an inflammatory stimulus, we hypothesized that PIAS1KO cells may be more sensitive to the pro-inflammatory signaling associated with HCMV infection. To examine this possibility, we analyzed the expression of the three ISGs examined above, IFIT3, BST2, and ISG15. Consistent with our previous results, there was no significant difference in the expression of these genes in either cell line under non-treated conditions (Fig. 4h-j). In contrast, either TNFα-treatment or ΔUL26 infection of PIAS1KO cells strongly induced the expression of these ISGs relative to control cells (Fig. 4h-j). In addition, the inactivation of PIAS1 also resulted in the strong induction of ISG expression during WT HCMV infection (Fig. 4h-j), suggesting that PIAS1 is important in limiting ISG expression during normal HCMV infection. During ΔUL26 infection, the expression of these ISGs was elevated, but to a lesser extent (Fig. 4h-j), suggesting that UL26 is necessary for PIAS1 to fully suppress ISG gene expression during infection. Taken together, these experiments indicate that inactivation of PIAS1 potentiates anti-viral transcriptional responses resulting in the strong induction of cytokine-signaling genes upon HCMV infection.

### PIAS1 inactivation sensitizes cells to cytokine treatment to limit viral infection

Given PIAS1’s role in modulating cytokine signaling-associated gene expression, we examined how it contributes to cytokine-mediated inhibition of HCMV infection. PIAS1KO cells were significantly more sensitive to TNFα and IFNα with respect to their ability to limit viral infection relative to ntg controls (Fig. 5a-b). In contrast, PIAS1 inactivation had little effect on IFNγ’s anti-viral activity (Fig. 5c). The ΔUL26 virus is sensitive to TNFα pre-treatment (17), and this sensitivity is exacerbated by the inactivation of PIAS1 (Fig. 5d), suggesting that in the face of TNFα treatment, the loss of PIAS1 compounds the cytokine-associated sensitivity of the ΔUL26 mutant. To target PIAS1 orthogonally, we utilized lentiviral-mediated CRISPRi to inhibit PIAS1 transcription (38, 39) (Fig. 5e). Cells expressing the more effective CRISPRi sg1 guide induced more significant anti-viral activity in response to TNFα and IFNα (Fig. 5e-f). Collectively, these experiments indicate that PIAS1 limits cytokine-induced anti-viral activities during HCMV infection.

**Figure 5.**
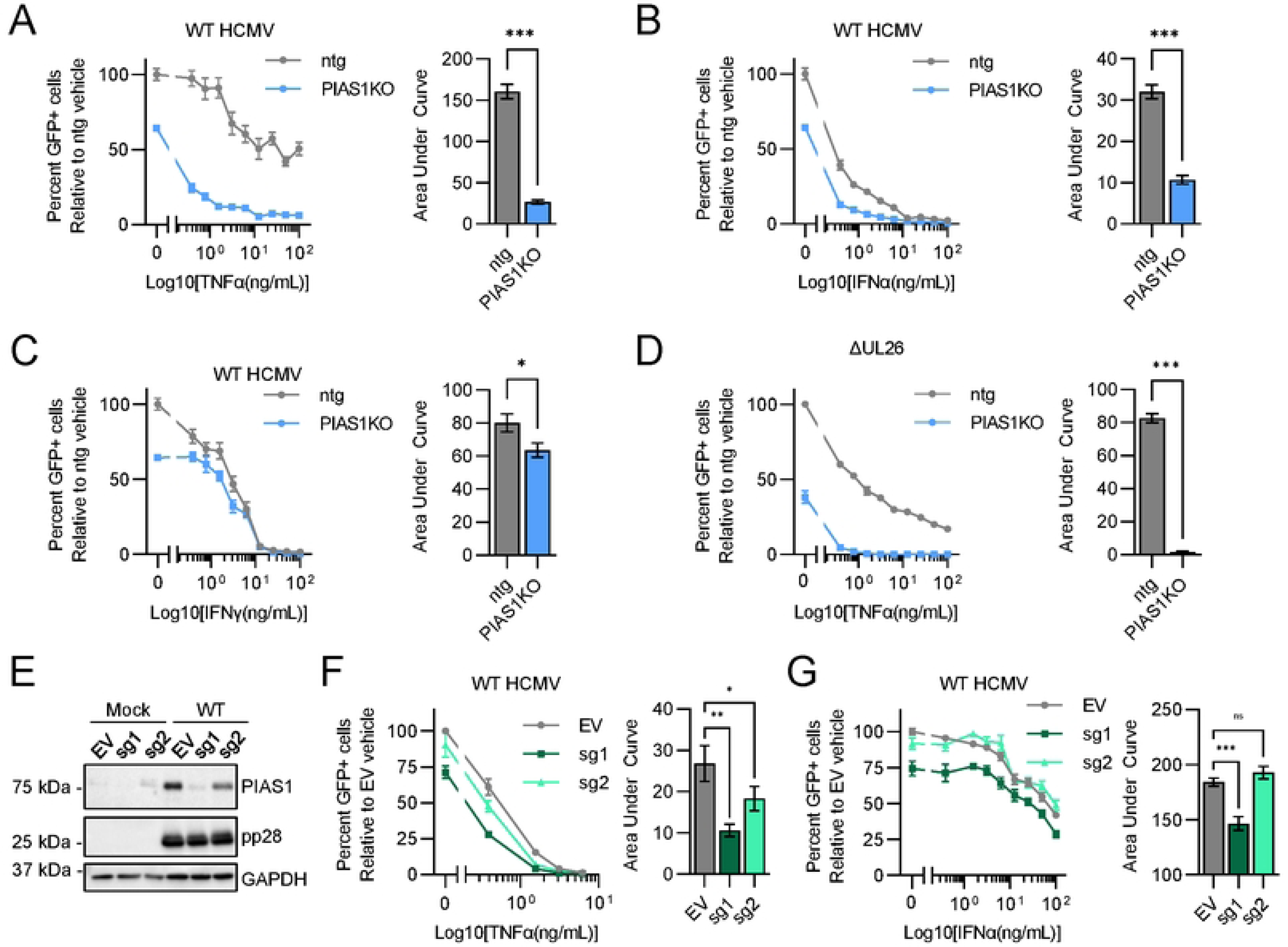
PIAS1 inactivation sensitizes cells to anti-viral cytokine treatment. (A - C) ntg (grey) and PIAS1KO (light blue) cells pre-treated with control medium or a 9-point dose curve of TNFα (A) IFNα (B) IFNγ (C) for 24 hr then infected with GFP-expressing wildtype HCMV (wtHCMV, MOI = 0.05). At 6 days post infection (dpi), total number of infected (GFP+) cells were quantified for each condition and normalized to control-treated ntg (100%). (D) ntg (grey) and PIAS1KO (light blue) cells pre-treated with control medium or a 9-point dose curve of TNFα for 24 hr then infected with GFP-expressing ΔUL26 (MOI = 0.1). At 7 days post infection (dpi), total number of infected (GFP+) cells were quantified for each condition and normalized to control-treated ntg (100%). (E and F) Cells transduced with CRISPRi plasmid containing empty vector (EV. grey) or PIAS1 sgRNAs (sg1, dark green & sg2, light green) constructs pre-treated with TNFα (E) or IFNα (F) for 24 hr then infected with GFP-expressing wildtype HCMV (wtHCMV, MOI = 0.05). At 7 days post infection (dpi), total number of infected (GFP+) cells were quantified for each condition and normalized to control-treated ntg (100%). Data is represented as the mean of 6 biological replicates (A - D) or 12 biological replicates (E and F) +/-SEM as an XY-plot (left). Control pre-treatment is represented in the left segment of the plot as x = 0. Area under the curve was calculated for the right segment of each XY-plot, excluding the x=0 control pre-treatment) and is represented as a bar graph (right). (A - D) t-values determined by student’s t-test n.s = not significant, *t<0.033, **t<0.002, ***t<0.001 (E and F) FDR-adjusted p-values determined using 2-way ANOVA followed by two-stage step-up method of Benjamini, Krieger and Yekutieli n.s = not significant, *p<0.033. **p<0.002, ***p<0.001.

### PIAS1 inactivation attenuates wildtype HCMV infection

Inactivation of PIAS1 reduced the number of infected cells in the absence of cytokine treatment (Fig. 5a-c). We next tested how PIAS1 contributes to HCMV viral spread. Inactivation of PIAS1 with either RNP or CRISPRi restricted WT HCMV spread (Fig. 6a & b, respectively). In contrast, ΔUL26 viral spread was not significantly impacted by the absence of PIAS1 (Fig. 6c), suggesting that the contribution of PIAS1 to infection depends on the presence of UL26. Plaque size and the initiation of WT HCMV infection were also both reduced in PIAS1KO cells compared to ntg cells (>50% reduction, Fig. 6d-f). PIAS1 inactivation also reduced the plaque size and initiation of infection for ΔUL26 but to a lesser extent (∼25%, Fig. 6d, g & h). Viral spread of WT HCMV and TB40/e, a more clinically relevant strain of HCMV, was also restricted in PIAS1KO cells (Fig. 6i & j). Collectively, our data indicate that PIAS1 is important for HCMV infection and that these contributions depend in part on the presence of UL26.

**Figure 6.**
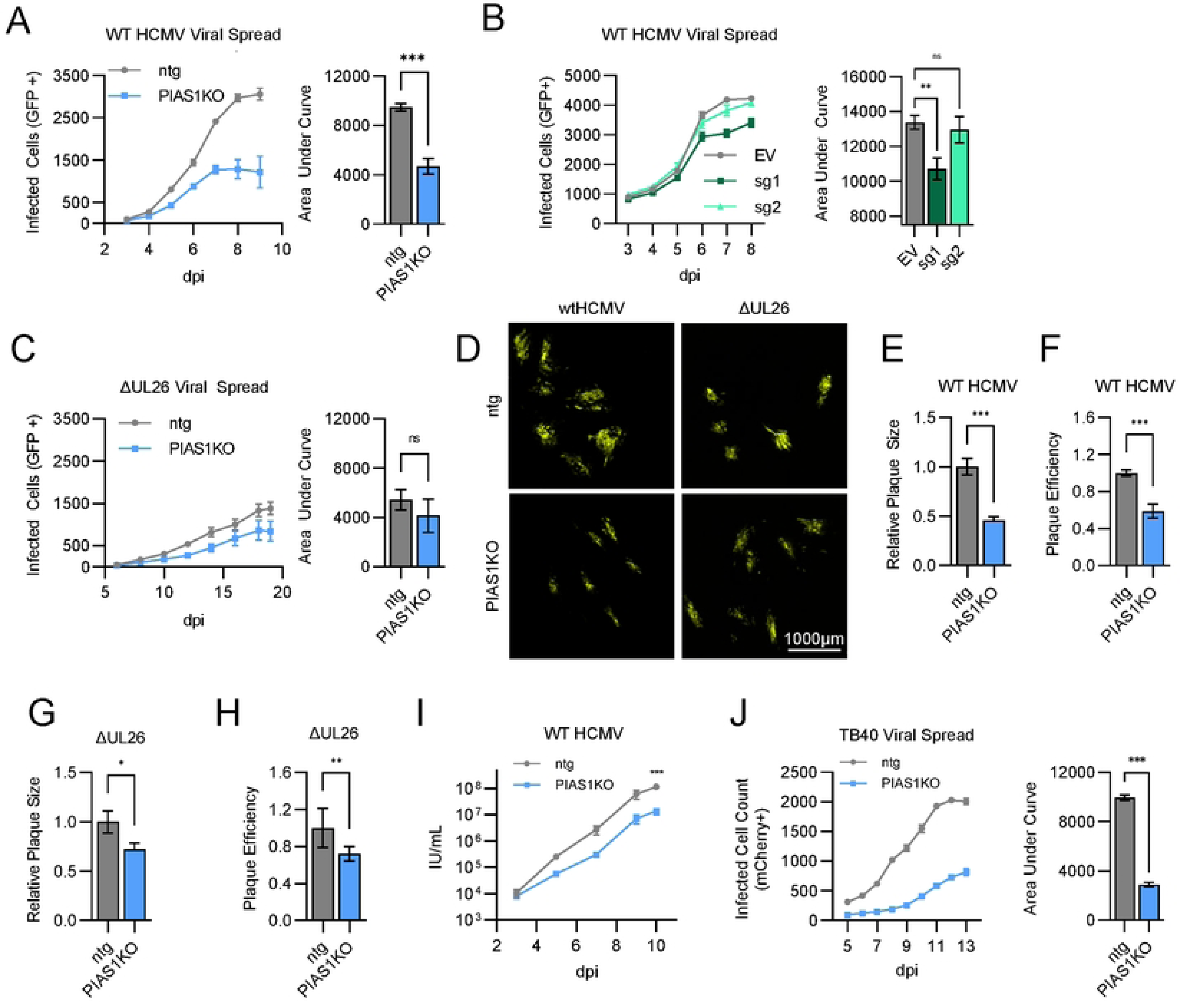
PIAS1 inactivation attenuates wildtype HCMV infection to a greater extent than ΔUL26 infection. (A) Viral spread of wildtype HCMV (wtHCMV) that expresses GFP at MOI = 0.01 in ntg (grey) and PIAS1KO (light blue) cells. Data represented as the mean of 6 biological replicates +/- SEM in an XY graph (left). Total area under curve (AUC) calculated for viral spread in each cell line and plot as a bar graph +/- SE (right). T-values determined student’s t-test n.s = not significant. *t<0.033. **t<0.002. ***t<0.001. (B) Viral spread of wtHCMV that expresses GFP at MOI = 0.05 in MRC5 cells transduced with CRISPRi plasmids containing empty vector (EV) or sgRNA targeting PIAS1 (sg1 dark green. sg2 light green). Data represented as the mean of 6 biological replicates +/- SEM in an XY graph (left). Total area under curve (AUC) calculated for viral spread in each cell line and plot as a bar graph +/- SE (right). FDR-adjusted p-values determined using 2-way ANOVA followed by two-stage step-up method of Benjamini, Krieger and Yekutieli n.s = not significant, *p<0.033. **p<0.002. ***p<0.001. (C) Viral spread of ΔUL26 that expresses GFP at MOI = 0 01 in ntg (grey) and PIAS1KO (light blue) cells. Data represented as the mean of 6 biological replicates +/- SEM in an XY graph (left). Total area under curve (AUC) calculated for viral spread in each cell line and plot as a bar graph +/- SE (right). T-values determined student’s t-test n.s = not significant, *t<0.033, **t<0.002, ***t<0.001. (D - H) Ntg and PIAS1KO cells infected with a low number of plaque forming units of wtHCMV or ΔUL26 that expresses GFP. (D) Representative images of plaques at 8 dpi. (E and G) Plaque size data is represented as the mean area of 20 representative plaques from each group normalized to the mean plaque size of wtHCMV in ntg cells +/- SE. (F and H) Plaque efficiency data is represented as the mean plaque count from 4 independent biological replicates +/- SE. (E - H) T-values determined student’s t-test n.s = not significant, *t<0.033, **t<0.002, ***t<0.001. (I) Viral growth assay of wtHCMV in ntg and PIAS1KO cells. Cells infected with wtHCMV (MOI = 0.1) and medium harvested from infections at the indicated times post infection (dpi). Infectious units (IU) per mL was quantified for each sample and plotted as the mean +/- SEM of 4 biological replicates. T-values determined by student’s unpaired t-test, ns = not significant, *t<0.033, “t<0.002, **t<0.001. (J) Viral spread of the HCMV clinical strain TB40 that expresses mCherry at MOI =0.1 in ntg (grey) and PIAS1KO (light blue) cells. Infected cells (mCherry positive) quantified by live-cell imaging every 24 hr starting at 5 days post infection (dpi) for 13 days. Data represented as the mean of 6 biological replicates +/- SEM in an XY graph (left). Total area under curve (AUC) calculated for viral spread in each cell line and plot as a bar graph +/- SE (right). T-values determined student’s t-test n.s = not significant, *t<0.033, **t<0.002, ***t<0.001.

## Discussion

Innate immune evasion is an important determinant of successful viral replication. Viruses have developed strategies to evade detection and attenuate anti-viral signaling to support high titer replication. In this study, we find that the full-length UL26 protein proximally interacts with a complex composed of PIAS and STAT family members. Further, our data indicate that UL26 is necessary to broadly restrict STAT family member activation during infection and that UL26 is sufficient to attenuate cytokine-induced IFN and TNF signaling (Fig. 2). Our results show that similar to UL26, the cellular PIAS1 protein is critical for limiting anti-viral gene expression during HCMV infection and upon cytokine challenge. Inactivation of PIAS1 substantially attenuates WT HCMV infection but has a reduced anti-viral impact on ΔUL26 infection, suggesting that UL26 is necessary for HCMV to benefit from PIAS1’s activity. Collectively, our data suggest that the UL26-PIAS1 complex attenuates anti-viral signaling during HCMV infection.

To identify UL26 interacting proteins that were more likely to be relevant to UL26’s contributions to HCMV infection, we compared the interactomes between TurboID-tagged full-length UL26 and a non-functional UL26 mutant, containing a 38-amino acid C-terminal deletion (19). Of the 68 proteins preferentially biotinylated by full-length UL26, 67 were human proteins. Many of these were STAT and PIAS family members (Fig. 1), which are well-known modulators of anti-viral signaling, and these interactions are consistent with UL26’s role in modulating cellular anti-viral defense. In addition, one viral protein was preferentially biotinylated by full-length UL26, UL31, which was ∼8-fold enriched in the wildtype UL26 sample relative to the C-terminal deletion (Table S3). UL31 can inhibit anti-viral signaling via cGAS inhibition (40). This raises the possibility that UL26 and UL31 might cooperate to modulate anti-viral signaling during infection, although, for both UL26 and UL31, their expression in isolation is sufficient to attenuate anti-viral gene expression (40, 41).

Previously, a global analysis of HCMV protein interactomes included the identification of proteins that interacted with stably expressed UL26 during infection (42). The study identified 44 human proteins that putatively interact with UL26 (42), including two that we identified here, CREB-binding protein (CREBBP) and histone acetylase p300 (EP300). Both these proteins have been found to interact with multiple partners in the STAT-PIAS complex depicted in Fig. 1i. CREBBP and EP300 are intrinsically disordered proteins that modulate transcriptional activation, including the regulation of anti-viral gene expression (43, 44). While these two proteins were common between the earlier study and ours, the majority of the putative interactors were different in the two studies. This likely reflects the different timing of when the pulldown samples were processed. Here, we biotinylated proteins at twenty-four hours post-infection, a time at which UL26 localizes to the nucleus (15, 45), whereas the previous interactome study performed pull-downs at sixty hours post-infection, a substantially later point of the viral life cycle, and a time at which UL26 is largely cytoplasmic (15, 42, 45). These times reflect very different viral life cycle stages, and thus likely reflect differences in UL26 interactions, e.g., modulation of host gene expression in the nucleus versus viral packaging into the tegument that occurs in the cytoplasm. The previous interactome also identified five UL26-interacting viral proteins (42), one of which, UL25, was also identified in our proximity-labeling experiment. UL25 was biotinylated by wildtype UL26 and UL26ΔC to a similar extent, suggesting this interaction is not dependent on the presence of the C-terminus of UL26. Little is known about the function of UL25 during infection. It is a nonessential protein located in the tegument (14, 46), and is possibly involved in virion envelopment (47).

UL26 was found to interact with several PIAS and STAT proteins and, other innate immune modulators previously found to form complexes (Fig. 1i) (24). The known roles that these cellular proteins play in cellular immunity, coupled with UL26’s ability to block STAT and TNFα signaling strongly implicate these interactions as being important for HCMV-mediated inhibition of anti-viral defenses. However, it remains to be elucidated which proteins UL26 is physically binding, and how UL26 binding might affect their activity. Despite these questions, UL26 and PIAS proteins can broadly modulate both STAT and NFkB signaling (21, 48), suggesting that a UL26-PIAS protein interaction may limit anti-viral signaling during infection.

The PIAS family of proteins are SUMO-ligases that contain a RING-finger-like zinc-binding domain (RLD) essential for their SUMO-ligase activity and a highly acidic (AD) domain that facilitates SUMO binding (20, 49). SUMOylation is a dynamic post-translational modification that regulates a number of molecular functions, including chromatin structure, DNA repair, transcription, innate immune signaling, cell cycle progression, protein trafficking, and the anti-viral response (27, 48, 50). Notably, PIAS1 can SUMOylate STAT1 at Lys703, which inhibits STAT1’s DNA binding and transcriptional activity following IFNγ-stimulation (33, 51). Similarly, PIAS proteins can inhibit NFkB signaling via binding and/or SUMOylating NFkB transcription factors and upstream regulators (35, 52). Further, SUMO-1 over-expression has been found to inhibit NFκB-dependent transcription (53). Here, we find that treatment with a SUMOylation inhibitor Subasumstat (32), attenuates HCMV replication (Fig. 3e-g). These data suggest that SUMOylation is broadly important for HCMV replication, however, the pleiotropic effects associated with pharmacological inhibition of SUMOylation can make precise conclusions difficult as SUMO modifications have been found to have both pro- and anti-viral effects on HCMV infection (30). The HCMV IE1 and IE2 proteins are SUMOylated during infection (54, 55, 56, 57, 58), as is the viral DNA polymerase processivity factor, UL44 (59, 60). In various instances, these SUMO modifications have been found to both support and inhibit infection, suggesting that the modulation of SUMOylation activity and the impact on infection is nuanced and context-specific.

We find that UL26 is sufficient to block IFNα and TNFα induced transcriptional activity (Fig. 3), indicating that UL26 can broadly suppress anti-viral defenses. PIAS1 has exhibited similar activities, that is, inhibiting STAT and NFkB signaling (21, 35, 61). Further, in over-expression experiments in the absence of infection, PIAS1 has been found to SUMOylate the HCMV IE2 protein, although the functional consequences during infection were not explored (58). We found that the inactivation of PIAS1 on its own did not dramatically alter the expression of STAT or NFκB-responsive genes but instead modulated genes involved in cholesterol metabolism (Fig. 4), a connection that has been previously observed (36, 37). However, upon stimulation by either HCMV infection or TNFα, cytokine signaling genes were strongly induced, largely phenocopying ΔUL26 infection (Fig. 4), indicating that the lack of PIAS1 strongly potentiates anti-viral gene expression in the face of an inflammatory challenge. Consistent with this finding, the inactivation of PIAS1 made cells more resistant to viral infection, and more capable of limiting viral infection at lower doses of TNFα treatment (Fig. 6). Notably, while the inactivation of PIAS1 substantially curtailed the viral spread of WT HCMV, there was no significant change in the spread of the ΔUL26 virus (Fig. 6c). These results suggest that UL26 is necessary for HCMV to take advantage of the presence of PIAS1, consistent with a model in which a UL26-PIAS1 complex attenuates anti-viral gene expression.

The specific mechanisms through which UL26 and PIAS1 modulate anti-viral gene expression remain to be elucidated. PIAS1 can alter transcription factor activity via transcription factor SUMOylation, direct binding and inhibition of transcription factors in the absence of SUMOylation, and through direct DNA binding at promoter sites, resulting in transcriptional repression (20, 48). It remains to be determined if UL26 specifically induces any of these mechanisms. Further, while we focused primarily on PIAS1, there are three additional human PIAS proteins, PIAS2, PIAS3, and PIAS4, that were all biotinylated preferentially by full-length UL26 and that have immunomodulatory activities (20, 52, 62). It remains to be determined to what extent UL26 directly interacts with the various PIAS family members and modulates their activity.

Regulation of cellular inflammatory and anti-viral gene expression programs is a central host-pathogen interaction that governs host defense. Our results indicate that productive HCMV infection relies on the UL26 and PIAS1 to prevent activation of anti-viral gene expression during infection. In this regard, HCMV has hijacked the host cell’s natural anti-inflammatory control mechanisms to limit anti-viral responses. A better understanding of the mechanisms involved could provide novel insights into these critical pathways and potentially highlight novel avenues for anti-viral therapeutic intervention.

## Materials and Methods

### Tissue Culture and Virus Propagation

Telomerase-transduced MRC5 lung fibroblasts and HEK293T cells were cultured in Dulbecco’s modified Eagle serum (DMEM; Invitrogen #11965118) supplemented with 10% (v/v) fetal bovine serum (FBS), and 1% penicillin-streptomycin (Pen-Strep; Life Technologies #15140-122) at 37°C in a 5% (v/v) CO_2_ atmosphere.

All AD169 viral stocks were derived from the BAd wt clone of AD169 Genebank accession number: FJ527563. The WT strain of HCMV used in experiments from Bad wt AD169 is referred to in this paper as WT HCMV. The GFP expressing wildtype AD169 referred to as WT-GFP, was BADsubUL21.5 (14). The UL26-deletion virus is a GFP transposon insertion, herein referred to as ΔUL26 (14). The truncated mutant of UL26 where the 185^th^ codon was replaced with a stop codon is herein referred to as UL26ΔC (19). The TB40/E-mCherry virus was a generous gift from Christine M. O’Connor and Eain Murphy, generated as described here (63). Recombinant viruses created for this work are described below in ‘BAC Recombineering’. All viral stocks used in this manuscript were propagated in MRC5 cells and concentrated by ultracentrifugation. Briefly, MRC5s were grown to confluence and placed in serum-free DMEM for 24 hr. Cells were infected at MOI = 0.01 (or 0.1 for viruses with a known growth defect, e.g. UL26-mutants). Infected cells were scraped into the medium and transferred to a sterile 50 mL conical. Cells were collected by centrifugation at 3,000 RPM for 5 minutes at 4C and virus-containing supernatant was set aside in a sterile Erlenmeyer flask. The cell pellet was resuspended in 5 mL virus-containing supernatant and transferred to a polystyrene tube. The cell solution was vortexed for 30 seconds and sonicated for 30 seconds three times then flash-frozen on dry ice and thawed in a 37C water bath to lyse cells. Cell debris was collected by centrifugation at 3,000 RPM for 5 minutes at 4C. The supernatant was combined with virus-containing supernatant in the Erlenmeyer flask. The medium was mixed well and distributed equally among ultracentrifuge tubes. Sorbitol buffer (20% D-sorbitol, 50 mM Tris-HCl, 1 mM MgCl_2_) was underlaid in each ultracentrifuge tube and the virus was collected at 26,000 rpm for 1.5 hrs at 23C. The supernatant was removed from the virus pellet by aspiration and the virus was suspended in a 1/25 volume ratio of original culture volume of 50:50 filter-sterilized solution of PBS supplemented with 3% BSA and DMEM supplemented with 10% FBS. Stocks were titered using a modified Reed & Muench TCID50 calculator from the Lindenbach lab (64).

For experiments involving infection, MRC5 cells were grown to confluence and then placed in serum-free DMEM supplemented with 1% PenStrep for 24 hours prior to infection. For experiments where cells are treated with cytokines or drugs prior to infection, MRC5 cells were grown to confluence and then placed in serum-free DMEM supplemented with 1% PenStrep and the indicated treatment for 24 hours prior to infection unless otherwise stated. To infect, the medium on cells was replaced with the minimum volume of viral adsorption medium to cover the monolayer of cells containing the indicated MOI in serum-free DMEM supplemented with 1% PenStrep. After 2.5 hr adsorption, the medium was removed and replaced with fresh serum-free DMEM supplemented with 1% PenStrep unless otherwise stated. Cells harvested as indicated at the timepoint of interest as described below.

### Reagents and Preparation of Treatments

Human tumor necrosis factor alpha (TNFα, GoldBio #1130-01-100), Interferon alpha (IFNα, Krackler # 45-I4276-20UG), and Interferon gamma (IFNγ, Medchemexpress #HY-P70610-50UG) were solubilized to 100 μg/mL concentration in sterile H2O and stored as 12 μL aliquots at -80C. Subasumstat (SUB, Fisher Scientific #50-217-3396) was solubilized to 10 mM in DMSO and stored at 20 μL aliquots at -80C. Biotin (Sigma, #B4501-100MG) was stored as a powder at room temperature and solubilized in serum-free DMEM to the indicated concentration immediately before use.

### Construction of plasmids

Primers used to assemble the fusion construct of UL26 to TurboID and clone into pLenti CMV/TO expression plasmid (addgene, #22262) are listed in Table S10. The TurboID plasmid (3xHA-TurboID-NLS_pcDNA3, addgene #107171) and UL26 fragment were amplified to have homology arms to pLenti plasmid at the attb1/BamHI cut site and a 3x GGGGS linker repeat and purified. The pLenti vector was digested with BamHI (NEB #R0136) and XbaI (NEB #R0145) following the manufacturer’s instructions overnight. DNA fragments were combined with the digested pLenti plasmid in a Gibson Assembly reaction mix (NEB #M5510AA) at a molar ratio of 1:3:3 [vector:TurboID:UL26] for 1 hour at 50C. Following this reaction, STBL3 competent cells were transformed with 2 μL of the Gibson Assembly mixture and colonies were grown on LB agar plates supplemented with Ampicillin at 37C overnight. Colonies were selected for colony PCR using primers that flank the construct insert region (See Table S10, pLenti plasmid sequencing primers). DNA samples of the appropriate size were submitted to Azenta for Sanger Sequencing. Colonies containing plasmids with the appropriate sequence were grown in LB and stored in 50% glycerol at -80C.

### Bacterial Artificial Chromosome (BAC) Recombineering

Recombinant virus containing TurboID fused to UL26 and UL26delC in the UL26 ORF were created as previously described(19) using the Bad wt clone of AD169 Genebank accession # FJ527563. The two-step PCR method for recombineering was followed as previously described using a Bad wt clone of AD169 with Kan/Isce I cassette in the UL26 ORF housed in GS1783 cells which contain an arabinose-inducible I-Sce 1 restriction site used for negative selection(19, 65). TurboID fused to UL26 wildtype or UL26ΔC were amplified to have homology arms to regions flanking the UL26 ORF of Bad WT clone of AD169, now containing Kan/Isce I cassette (BAd169+Kan), primers used are listed in Table S10, BAC Recombineering Primers. Amplified DNA oligos containing TurboID fused to UL26 wildtype or UL26ΔC were then transformed into GS1783 electroshock competent cells containing BAd169+Kan and resulting colonies were negatively selected on LB +L-Arabinose plates in the presence or absence of Kanamycin. Colonies that were no longer Kan-resistant were selected for colony PCR using a Touchdown method with primers flanking the UL26 ORF. Amplified DNA was run on an ethidium bromide agarose gel to identify colonies with DNA fragments of the expected size. These fragments were submitted to Azenta for Sanger Sequencing.

### Propagation of Recombinant Virus

BAC DNA housing recombinant virus of interest was isolated by growing GS1783 containing BAC of interest in 10 mL LB medium overnight. Cells were harvested by centrifugation at 4,000 RPM for 10 minutes at 4C. The cell pellet was resuspended gently in 200 μL cold buffer P1 from a Qiagen miniprep kit (#27106). Resuspension incubated at room temperature for 5 minutes. 200 μL buffer P2 from a Qiagen miniprep kit was added to the cell resuspension and incubated in ice for 5 minutes. Following this incubation period, 300 μL buffer P3 from a Qiagen maxiprep kit (#12163) was added to the sample tube and placed on ice for 10 minutes. The sample was centrifuged at 14,000 RPM for 10 minutes and supernatant was collected in a 2 mL sterile microcentrifuge tube. One mL of room temperature isopropanol was added to precipitate DNA and the tube was inverted 3x to mix. The sample was centrifuged at 14,000 rpm for 10 minutes at 4C. The supernatant was discarded and the pellet was resuspended in 500 μL filter-sterilized TEN buffer (25 mM Tris pH 7.6, 1 mM EDTA, 150 mM NaCl). The sample was incubated at room temperature for 10 minutes and then centrifuged at 14,000 rpm for 30 seconds. The supernatant was transferred to a clean 1.5 mL microcentrifuge tube and 1 mL prechilled 100% ethanol was added to the supernatant. The sample was inverted 3x and centrifuged at 14,000 RPM for 10 minutes at 4C. The supernatant was discarded and open tubes were placed in a sterile hood for 5 minutes for ethanol to evaporate. DNA was suspended in 50 μL sterile double-deionized water.

MRC5s were transfected with isolated BAC DNA. At 70-80% confluence, MRC5 fibroblasts were trypsinized and cells were isolated by centrifugation at 1.5 rpm for 5 minutes at 4C. The supernatant was discarded and cells were resuspended in 500 μL solution 6 (140 mM Na phosphate buffer, 5 mM KCl, 10 mM MgCl2, pH ∼7.2). 25 μL BAC DNA prep and 1 ug pp71 expression plasmid was added to the cell suspension. The sample was transferred to a 4 mm electroporation cuvette and cells were electroporated using BioRad Gene Pulser Xcell total system (#1652266) set to exponential, 265 V, 960 μF with a time constant between 15 and 20. Immediately after transfection, the electroporation mixture was added to a sterile 10 cm tissue culture dish containing 8 mL DMEM supplemented with 10% FBS and 10 mM Na Butyrate. Cells were placed in a 37C incubator for 2 hours to settle, then the medium was aspirated and replaced with fresh DMEM supplemented with 10% FBS. Cells were placed in a 37C incubator overnight and monitored every day until confluent. Once confluent, cells were transferred to a 15 cm tissue culture dish by trypsinization and grown to confluence. Cells were monitored every day for signs of infection and the virus was harvested from cells 2 days after 100% cytopathic effect (CPE) was apparent as described above.

### Proximity Proteomics Sample Preparation and Analysis

For each condition, approximately 1e7 MRC5 cells were grown to confluence and placed in serum-free DMEM medium for 24 hours. Cells were mock infected or infected with TurboID-N-UL26wt or TurboID-N-UL26ΔC (MOI = 3) using 9 mL of viral adsorption medium for 2.5 hours. The adsorption medium was removed and replaced with serum-free DMEM. At 23 hours post-infection, the medium was replaced with serum-free DMEM supplemented with 1 μM Biotin. After 1 hour of incubation with biotin, the medium was removed and cells were washed 2x with cold PBS. Cells were harvested in 5 mL RIPA lysis buffer (50 mM Tris pH 7.4, 150 uM NaCl, 1 mM EDTA, 0.25% (wt/v) Sodium Deoxycholate, 1% (v/v) TritonX-100, and 0.1% SDS) and transferred into a chilled 15 mL centrifuge tube, vortexed for 30 seconds and placed on ice for 20 minutes. Lysates were vortexed for 30 seconds and centrifuged for 15 minutes at 14,000 rpm at 4C.

To prepare beads for affinity pulldown, 100 μL streptavidin-bound magnetic beads (Dynabeads™ MyOne™ Streptavidin T1, ThermoFisher #A1254302) per sample were washed with 5x volume RIPA without detergent 3 times for 5 minutes each wash. Beads were resuspended in the original volume and kept on ice until ready to use.

Samples were incubated with 100 μL prepared magnetic beads for 1 hour at room temperature. The supernatant was removed from each sample and beads were washed with 1 mL Wash Buffer 1 (2% SDS in H2O) for 5 minutes on a rotator. The supernatant was removed and beads were washed with 1 mL Wash Buffer 2 (50 mM HEPES, 500 mM NaCl, 1 mM EDTA, 0.1% (v/v) TritonX-100) for 5 minutes on a rotator. The supernatant was removed, and beads were washed with 1 mL Wash Buffer 3 (10 mM Tris-Cl pH 7.4, 250 mM LiCl, 1 mM EDTA, 0.1% (wt/v) sodium deoxycholate) for 5 minutes on a rotator. The supernatant was removed and beads were washed with 1 mL 50 mM Tris (pH 7.4) for 5 minutes on a rotator. The supernatant was removed and protein was eluted from the beads with 40 μL Disruption Buffer (2% SDS, 50 mM Tris pH 7, 5% β-mercaptoethanol, 2.5% (v/v) Glycerol, Bromophenol Blue). Samples were boiled in Disruption Buffer for 5 minutes and centrifuged at 14,000 RPM for 10 minutes. The supernatant was transferred into a new Eppendorf tube, 10 μL affinity pulldown sample was run on a 10% acrylamide SDS-PAGE gel at 115 V for 1.5 hours. This gel was stained with Pierce silver stain for mass spectrometry (#B4501). The remaining 30 μL sample was run on a 10% acrylamide SDS-PAGE gel at 100 V for 15 minutes and stained with Coomassie (Bio-Rad #P1610786) following the manufacturer’s instructions. The gel was brought to the University of Rochester Proteomics core for sample prep and analysis.

The sum of the peak areas for each peptide that could be integrated for each poritein were represented at abundance for each protein. Samples with non-detected abundance values were replaced with the minimum abundance detected for the data set (1.03e4, 912 values replaced). Each abundance value was normalized to the median abundance value for that sample and multiplied by 1e7. Normalized values for Tur-N-UL26WT +Biotin and Tur-N-UL26ΔC +Biotin were graphed together as a scatter plot (Fig. 4.1d) with a line of identity (m=1).

Human proteins 5-fold enriched in Tur-N-UL26WT+Biotin relative to Tur-N-UL26ΔC (67) were uploaded to the Human Reference Protein Interactome Mapping Project (query-query)(24) to identify an interactome of 12 proteins that were preferentially biotinylated by Tur-N-UL26WT. This data was used to create Fig. 1i. The list of Human proteins 5-fold enriched in Tur-N-UL26WT+Biotin relative to Tur-N-UL26ΔC (67) was also analyzed by reactome.org Analysis Tool (66). The gene list was projected to human identifiers to provide the data in Table S2. Entities FDR values were -log_10_ transformed to provide the sata in Fig. 3a.

### CRISPR Ribonucleoprotein (RNP)-Based Knockout

Gene disruption by CRISPR-Cas9 ribonucleoprotein (RNP) was performed with the Neon^TM^ Transfection System 10 μL kit (ThermoFisher #MPK1025) as described previously(67) with electroporation settings voltage: 1100 V, width: 30 ms, pulses: 1. Guide RNAs were ordered from Synthego:

Negative Control Scrambled sgRNA (modified) #1: 5’-GCACUACCAGAGCUAACUCA-3’

RELA (sgRNA EZ kit): 5’-UCCCCUGGCAGAGCCAGCCC-3’

STAT1 (Gene Knockout Kit v2): sgRNA1: 5’-AGUGGUUAGAAAAGCAAGAC-3’; sgRNA2: 5’-AAAGCUGGUGAACCUGCUCC-3’; sgRNA3: 5’-UUCCCUAUAGGAUGUCUCAG-3’

PIAS1 (Gene Knockout Kit v2): sgRNA1: 5’-GCCAGCCUUUAGCAAAUGCA-3’; sgRNA2: 5’-CAGGCGUCAUGAUUUUCUGU-3’; sgRNA3: 5’-GGAGACAAAGUUGCUGGCAU-3’

### CRIPSRi Gene Knockdown

Two independent guide RNAs directed to the PIAS1 gene locus (sg 5’-CGGACAGTGCGGAACTAA-3’, sg2 5’-TAGGGAGTCCGGAGGTAG-3’) were designed with a 5’ and 3’ flanking sequence to a CRISPRi Lentivirus (LV) PURO plasmid (pLV hU6-sgRNA hUbC-dCas9-KRAB-T2a-puro with stuffer), gifted by the Harris Lab at University of Rochester Department of Biomedical Genetics (5’-GGAAAGGACGAAACACCG-3’ & 5’-GTTTTAGAGCTAGAAATAGC-3’, respectively). Guides with flanking sequences to the CRISPRi LV plasmid were amplified by PCR, cloned into the CRISPRi plasmid by Gibson Assembly as described above, and transformed into STBL3. Plasmids were isolated from positive colonies were screened by Sanger Sequencing to verify the PIAS1 sgRNA sequence. Lentivirus propagated in HEK293T cells and used to transduce MRC5s as previously described(16).

### Western Blotting

Protein detection was performed as previously described(68). The following antibodies were used for western blot analysis following the manufacturer’s instructions: STAT1 (Cell signaling, #D1K9Y), Phospho-STAT1 (Cell Signaling, #58D6), STAT2 (Cell Signaling, #D9J7L), Phospho-STAT2 (Cell Signaling, #D3P2P), GAPDH (Cell Signaling #5174), STAT3 (Cell Signaling #79D7), Phospho-STAT3(Tyr705) (Cell Signaling #D3A7), HA-Tag (Cell Signaling #2367), PIAS1 (Cell Signaling #D33A7), PIAS2 (ABclonal, #A5654), PIAS3 (Cell Signaling, #9042), PIAS4 (Cell Signaling, #4392), SUMO1 (Cell Signaling, #4930S), SUMO2/3 (Cell Signaling, #4971S), viral proteins IE1(69), pp28(70), UL26(15) (Mouse mAb).

For the detection of biotinylated proteins, cell lysates were run on a gradient SDS-PAGE gel and transferred onto a nitrocellulose membrane as described above. The membrane was then blocked in 1X TBST supplemented with 1% BSA and 0.2% TritonX100 for 10 minutes. The membrane was incubated in a diluted streptavidin conjugated to HRP (Fisher, PI-21130) solution (1:40,000 in 1xTBST supplemented with 1% BSA) for 40 minutes. Following this incubation, the membrane was washed 3x with 15 minutes with 1xTBST and imaged as described above.

For quantification of western blots, band intensities for GAPDH and protein of interest were determined using Image Lab v4.1 for each lysate. GAPDH intensity for all samples was divided by the maximum GAPDH intensity. Band intensity of the protein of interest was then normalized to the calculated GAPDH value and expressed as a percentage.

### Luciferase Assay

The ISRE firefly luciferase reporter plasmid was a gift from the lab of Dr. Ruth Serra-Moreno. HEK293T cells were seeded in white-walled 96 well plates. At 30% confluence, cells were treated with 1 μg total DNA (250 ng firefly expression plasmid, 250 ng renilla expression plasmid, 500 ng EV or UL26) and 1.2 μL Fugene transfection reagent. At 48 hr, treatments (IFNα or TNFα) were added to the appropriate samples. 24 hours later, luciferase assays were carried out as previously described(17) using a Promega dual luciferase reporter assay.

### Virion Production by TCID50

Cells were treated and infected as described above. At the desired time post-infection, the medium was removed from the cell monolayer, frozen on dry ice in cryovials, and stored at -80C. Samples were thawed in a 37C water bath, vortexed, and then centrifuged for 5 minutes at 3,000 RPM to pellet cellular debris. Virus-containing supernatants were serially diluted (1:10) eight times in a deep 96-well plate. Using a multichannel pipette, 50 μL of each dilution was transferred to each well within a row of a 96-well plate containing sub-confluent MRC5 cells in 50 μL growth medium (n=12). After 2-3 days post-infection, 100 μL serum-free DMEM was added to each well. Infected wells were identified 10 days post-infection and sample titers were calculated using a modified Reed & Muench TCID50 calculator from the Lindenbach lab(64).

### Plaque Efficiency and Size Assay

Plaque assays were performed in the indicated cell type as previously described(67). At 8-10 dpi, plaques were imaged using a Cytation 5 imaging reader (BioTek). Field of view (FOVs) with plaques were manually captured using a 4x magnification objective lens and predefined YFP channel with an excitation wavelength of 469 nm and emission wavelength of 525 nm for GFP+ cells. For each condition, at least 20 independent plaques were captured. Plaque-size objects within each FOV were defined with Gen 5 Image Primer 3.11 software on the YFP image as objects at least 100 um in size and intensity threshold of 5,000 units, touching objects were not split.

### Viral Spread

MRC5s were grown to confluence in 96-well TC-treated black-walled plates and treated/infected as indicated. To detect infected (GFP+) cells over time, plates were imaged using Cytation 5 imaging reader (BioTek) using a 4x magnification objective lens and predefined YFP channel with an excitation wavelength of 469 nm and emission wavelength of 525 nm. Infected (GFP+) cells were defined with Image Primer 3.11 software on the YFP image as objects 20-200 um in size with an intensity threshold of 5,000 units, touching objects were split. For cytokine pre-treatment assays, GFP+ cells were quantified at 6 dpi for WT HCMV and 7 dpi for UL26TI. All samples were normalized to the number of infected (GFP+) cells in the untreated ntg control condition and expressed as a percent of 100.

### Viral Replication (Infectious Units/mL)

MRC5 cells were seeded into a 384-well black-walled plate (45 uL) and grown to confluence. Virus samples were loaded into rows A-D of a 96-well plate. Using an Integra Assist Plus instrument and 12-channel integra electric pipette, 5 μL of each virus sample from the 96 well plate was transferred to the 384 well plate and serially diluted 1:10 four times.

At 48 hpi, the number of infected cells in each well was determined using Cytation 5 imaging reader (BioTek) using a 4x magnification objective lens and predefined YFP channel with an excitation wavelength of 469 nm and emission wavelength of 525 nm. Infected (GFP+) cells were defined with Image Primer 3.11 software on the YFP image as objects 20-200 um in size with an intensity threshold of 5,000 units, touching objects were split. The total number of infected cells was considered ‘infectious units’ and the total infectious units/mL for each sample was calculated from the appropriate dilution.

### Click-Seq Library Construction and data analysis

First, 5 µg total RNA was subjected to poly(A) mRNA isolation according to the manufacturer’s protocols (NEB, #E7490). Then, 10 µL of the eluted mRNA was mixed with 1 µL 100 µM Genomic Adaptor_6N primer (5’-GTGACTGGAGTTCAGACGTGTGCTCTTCCGATCTNNNNNN-3’) and 2 µL of 5 mM AzNTP (Baseclick, #BCT-25∼ -28)/dNTP mixture in a 1:35 ratio. The mixture was then heated at 65°C/5min and snap-cooled on ice for 3min. Then, the reverse transcription was carried out using SuperScript III (ThermoFisher, #18080093). In brief, 7 µL master mix containing 4 µL 5X Superscript First Strand Buffer, 1 µL 0.1 M DTT, 1 µL RNaseOUT (ThermoFisher, #10777019), and 1 µL Superscript III was mixed with the RNA sample. Heating was performed as follows: 25°C/10 min, 50°C/40 min, and 75°C/15 min sequentially in thermocycler followed by RNase H (ThermoFisher, #AM2293) treatment using 1 U per reaction for 37°C/30 min and then 80°C/10 min. The cDNA was then purified using Sera-Mag Speedbeads (Cytiva, #65152105050250). Speedbeads working solution was made by washing 1 mL beads slurry twice in 1XTE buffer and then resuspending in 50 mL of 1XTE buffer containing 9 g PEG-8000, 1 M NaCl, and 0.05% Tween-20. Speedbeads, 1.8x reaction volume, was mixed with cDNA followed by a 5 min incubation at room temperature. The beads were then pelleted using a magnetic bead collector and the supernatant was discarded. Two washes with 80% ethanol were performed while the beads were pelleted on the magnet. The beads were then dried, and the cDNA was eluted by resuspending the beads in 22 µL 50 mM HEPES pH 7.2 for 2 min at room temperature. Then, 20 µL of the cDNA was mixed with 11 µL Click Mix and 4 µL of 5 µM UMI-click-adapter (5’-Hexynyl-NNNNNNNNNNNNAGATCGGAAGAGCGTCGTGTAGGGAAAGAGTGT-3’, HPLC purified). To activate the reaction, 4 µL of 50 mM vitamin C and 1 µL of Click Catalyst were pre-mixed before being added into the cDNA solution for 60min room temperature incubation in the dark. Clicked cDNA was then purified in the same manner as post-RT purification except eluted in 22 µL of 10 mM Tris pH 7.4. Then, half of eluted cDNA was subjected to PCR amplification with indexing primers i5 (5’-AATGATACGGCGACCACCGAGATCTACAC[Index]ACACTCTTTCCCTACACGACGCTCTTCC GATC*T-3’) and i7 (5’-CAAGCAGAAGACGGCATACGAGAT[Index]GTGACTGGAGTTCAGACGTGTGCTCTTCCGAT *C-3’). Both primers have a phosphorothioate bond added at the 3’ end (denoted as *) to increase stability. The PCR reaction was carried out in the volume of 50 µL including 25 µL 2X OneTaq Master Mix (NEB, #M0482), 2 µL of 5 µM i5 and i7 each, and cDNA in H_2_O with a total of 15 cycles of amplification given the 5 µg input RNA. The program was run at 94°C/4 min; 53°C/30 sec; 68°C/10 min; 15X [94°C/30 sec, 53°C/30 sec; 68°C/2 min]; 68°C/5 min; hold at 4°C. The library was then purified and size-selected with Speedbeads. In brief, 45 µL beads (0.9X volume of PCR reaction) were mixed with the reaction to remove larger fragments. The supernatant was collected and mixed with 10 µL beads (0.2X volume of PCR reaction) to remove smaller fragments and primer dimers. Two washes with 100% ethanol were then performed while the beads were on the magnet. Beads were dried before being resuspended in 12 µL of 10 mM Tris pH 7.4. The library was then collected and subjected to quality control on Agilent 2200 TapeStation and to analysis on Illumina NovaSeq 6000 at Genomics Research Core at the University of Rochester. Data was reported using Multi v 1.11 (71). Data was formatted with bcltofastq-2.19.1 and cleaned with fastp 0.23.1. Sequence reads were mapped to the human reference genome (GRCh38 + Gencode-38 annotation) using STAR_2.7.9a.

Protein coding transcripts were isolated from the data for further analysis (Table S5 & S6). For genes with multiple emsemble transcript accession numbers, the sample with the largest base mean value was selected to represent that gene. Genes with Log2 Fold Change > 2 and padj < 0.5 were analyzed by reactome analysis tool as described above (66) to provide the data in Table S6 & S7 and Figures 4b - f. Previously reported data (GSE137065) was re-analyzed using the same methods (see Table S8 & Fig. 3g). Data from this work (Table S7) was compared to previously reported data (GSE137065, Table S8) in Table S9. Intersecting genes were identified to produce the data represented in Fig. 4d & e.

### Analysis of RNA (RT-qPCR)

RNA was isolated from cells by Trizol extraction and used to synthesize cDNA with a qScript cDNA synthesis kit (Quantabio). Samples were analyzed by Real-Time qPCR and relative quantities of gene expression were calculated as previously described(72). The following primers were used:

BST2 F 5’-AACACGGTTAGCGGGGAGAG-3’; BST2 R 5’-GGGACTCATTGTCCGGAGGG-3’; ISG15 F 5’-CGGGCAACGAATTCCAGGTG-3’; ISG15 R 5’-CACGCCGATCTTCTGGGTGA-3’; IFIT3; F 5’-GGAAACAGCCATCATGAGTGAGG-3’; IFIT3 R 5’-TGAATAAGTTCCAGGTGAAATGGC-3’; GAPDH F 5’-CATGTTCGTCATGGGTGTGAACCA-3’; GAPDH R 5’-ATGGCATGGACTGTGGTCATGAGT-3’

### Statistics

All statistics were carried out using GraphPad Prism v10.0.1 unless otherwise indicated.

## Supporting Information

**S1 Table. Proximity proteomics data.** Proteomics data from biotin-enriched samples harvested as outlined in Fig. 1F. Description of columns: <u>Protein FDR Confidence</u>: The confidence that the software has that the given protein is present in the sample. In most cases we filter the data to show only high confident proteins. <u>Accession:</u> Uniprot or NCBI accession number. <u>Description:</u> Name of the protein, along with the species and gene name. The gene name can be found after “GN=”. <u>Marked As:</u> This column is present if multiple species were searched. It will show which database the given protein originated from. <u>MW [kDa]:</u> Molecular weight of the protein. <u># of Unique Peptides</u>: Peptides can be shared between different proteins or protein isoforms, so this column shows the number of peptides that are unique to the given protein group. <u># of Peptides</u>: The total number of unique and shared peptides for the given protein group. <u># of PSMs (Peptide Spectral Matches):</u> Peptide spectral matches are MS2 scans that are matched to a peptide by the search algorithm. This differs from the number of peptides because the same peptide can be fragmented and identified multiple times within a mass spec run, especially if the protein that the peptide originated from was in high abundance. The number of PSMs will therefore always be greater than or equal to the number of peptides identified. <u>Abundance Ratio (log2):</u> The protein fold change in log2 format between the given samples or groups. The software caps the maximum fold change at 100, so the largest log2 fold change you will see is 6.64 (2^6.64 =100). <u>Abundance Counts:</u> The number of peptides that are used to calculate the abundance of the given protein. <u>Abundances (or Normalized Abundances):</u> The sum of the peak areas for each peptide that could be integrated for the given protein. If normalization was selected, then the normalized abundances will be shown

**S2 Table. Normalized proximity proteomics data.** Median normalized protein abundance, from S1 Table, supplied data for Fig. 1h. Proteins 5-fold enriched in WT+B relative to UL26ΔC+B (dark green, right).

**S3 Table. HCMV proteins detected in proximity proteomics experiment.** HCMV proteins from S2 Table. Proteins 5-fold enriched in WT+B relative to UL26ΔC+B (dark green, right).

**S4 Table. Reactome analysis of genes enriched in WT UL26 sample.** Reactome analysis on the list of proteins 5-fold enriched in WT UL26 compared to UL26ΔC from proximity proteomics analysis, input list is from Table S2 represented in green (left). Reactome analysis report (right). The top 10 pathways scored by -log_10_(Entities FDR) are highlighted in orange and are represented in Fig. 3A.

**S5 Table. RNA-Seq data comparing vehicle treated PIAS1KO and ntg cells.** (A) RNA-Seq analysis of [PIAS1KO_Veh/ntg_Veh]. (B) Protein coding mRNA from A. (C) Protein coding mRNA with padj<0.05 from B. (D) Protein coding mRNA from C with Log_2_[PIAS1KO_Veh/ntg_Veh] > 2. (E) Reactome analysis of genes in D, pathways highlighted in blue are represented in Fig. 4B. This table also provides the data for Fig. 4C.

**S6 Table. RNA-Seq data comparing TNFα treated PIAS1KO and ntg cells.** (A) RNA-Seq analysis of [PIAS1KO_TNFα/ntg_TNFα]. (B) Protein coding mRNA from A. (C) Protein coding mRNA with padj<0.05 from B. This table provides the data for Fig. 4F.

**S7 Table. Reactome analysis of RNA-Seq data comparing TNFα treated PIAS1KO and ntg cells.** Genes significantly enriched in [PIAS1KO_TNFα/ntg_TNFα] from Table S6 (padj<0.05 and Log2FoldChange>2). Gene list (left) imported in to reactome analysis tool, reactome report (right). Reactome pathways highlighted in yellow are represented in Fig. 4D.

**S8 Table. Analysis of RNA-Seq data from GSE137065, cells infected with HCMV or ΔUL26 mutant.** (A) Genes depleted in cells infected with HCMV relative to ΔUL26 mutant virus, provides the data for Fig. 4F. (B) Genes significantly enriched in cells infected with ΔUL26 (Log_2_[HCMV/ΔUL26] < -2 & padj < 0.05). This gene list is the input for reactome analysis. (C) Reactome analysis report of B. Intersecting reactome pathways from top 10 reactome terms of [PIAS1KO+TNFα/ntg+TNFα] from Table S7 also significantly perturbed by the absence of UL26 are represented in yellow. The -log10(entities FDR) of these intersecting reactome pathways are represented in Fig. 4D.

**S9 Table. Identifying the genes enriched in the absence of UL26 or PIAS1.** Intersection of entities found in reactome terms shared by [PIAS1KO+TNFα/ntg+TNFα] from Table S7 and [HCMV/ΔUL26] from Table S8. Data for Fig. 4E-G.

**S10 Table. Primers.** Primers used for cloning, BAC recombineering, sequencing, and RT-qPCR.

